# A population of gray matter oligodendrocytes directly associates with the vasculature

**DOI:** 10.1101/2022.04.11.487873

**Authors:** Justine S. C. Palhol, Maddalena Balia, Fernando Sánchez-Román Terán, Mélody Labarchède, Etienne Gontier, Arne Battefeld

## Abstract

Oligodendrocyte lineage cells interact with the vasculature in the gray matter. Physical and functional interactions between blood vessels and oligodendrocyte precursor cells play an essential role in both the developing and adult brain. Oligodendrocyte precursor cells have been shown to migrate along the vasculature and subsequently detach from it during their differentiation to oligodendrocytes. However, the association of mature oligodendrocytes with blood vessels has been noted since the discovery of this glial cell type almost a century ago and this interaction remains poorly explored. Here, we systematically investigated the extent of mature oligodendrocyte interaction with the vasculature in mouse. We found that ~17% of oligodendrocytes were in contact with blood vessels in cortex, hippocampus and cerebellum. Contacts were made mainly with capillaries and sparsely with larger arterioles or venules. By combining light and serial electron microscopy, we demonstrated that oligodendrocytes are in direct contact with the vascular basement membrane, raising the possibility of direct signaling pathways and metabolite exchange with endothelial cells. During experimental remyelination in the adult, oligodendrocytes were regenerated and associated with blood vessels in the same proportion compared to control cortex, suggesting a homeostatic regulation of the vasculature-associated oligodendrocyte population. Based on the frequent and close association with blood vessels, we propose that vasculature-associated oligodendrocytes should be considered as an integral part of the vasculature microenvironment. This particular location could underlie specific functions of vasculature-associated oligodendrocytes, while contributing to the vulnerability of mature oligodendrocytes in neurological diseases.

## Introduction

Oligodendrocytes are known to contribute to neuronal plasticity through adaptive myelination and refinement of existing circuits (Fields et al., 2015; Hill et al., 2018; Hughes et al., 2018; McKenzie et al., 2014; Steadman et al., 2019). Oligodendrogenesis starts during embryonic brain development in the white matter and continues into adulthood in neocortical areas (Fard et al., 2017; Hill et al., 2018; Hughes et al., 2018; Vincze et al., 2008). Once formed, mature oligodendrocytes maintain their location and are long-lived cells (Tripathi et al., 2017; Yeung et al., 2014).

Interactions between oligodendrocyte lineage cells and the brain vasculature have been previously identified. During embryonic development, oligodendrocyte precursor cells (OPCs) migrate along the vasculature scaffold (Tsai et al., 2016), which can be also be observed in adult remyelination (Niu et al., 2019). On one hand, endothelial cell secreted transforming growth factor beta 1 (TGF-β1), brain-derived neurotrophic factor (BDNF) and fibroblast growth factor (FGF) influence differentiation and survival of OPCs (Arai and Lo, 2009; Paredes et al., 2021). On the other hand, OPCs can stimulate angiogenesis during development and postnatal myelination through an HIF-activated secretion of Wnt and vascular endothelial growth factor (VEGF), presumably to adapt local blood supply to the metabolic needs during OPC differentiation (Yuen et al., 2014; Zhang et al., 2020). OPCs also participate in maintaining the blood-brain barrier integrity by secreting TGF-β1, which increases tight-junction protein expression in endothelial cells (Seo et al. 2014). Moreover, in pathological tissue from multiple sclerosis patients, as well as in a mouse model, blood-brain barrier disruption occurs as consequence of aberrant OPC clustering around blood vessels (Niu et al., 2019), further suggesting that vasculature-OPC interaction is tightly regulated.

Similarly, the existence of functional interaction between mature oligodendrocytes and the vasculature has been established. Endothelial cell released endothelin activates g-protein coupled endothelin receptor B on mature oligodendrocytes, subsequently stimulating myelin formation in the pre-frontal gray matter (Swire et al., 2019). Moreover, a relationship between mature oligodendrocytes and the vasculature has been noted since the first description of this cell type (Del Río Hortega 1928). Vasculature-associated oligodendrocyte were found in white and gray matter of different species and a role in blood flow control by these oligodendrocytes was proposed (Cammermeyer, 1960). Additionally, associations of oligodendrocytes with blood vessels have been described in the hippocampus (Vinet et al., 2010). More evidence that oligodendrocytes associate tightly with the vasculature emerged, after a cluster of presumably vessel-associated oligodendrocytes was identified in a single cell sequencing dataset analyzing zonation of the brain vasculature (Vanlandewijck et al., 2018). However, surprisingly little is known about the underlying structural properties of the interaction of mature oligodendrocytes with the vasculature in the adult neocortex.

In this study, we investigated the arrangement and frequency of oligodendrocyte-vasculature interaction in the neocortical gray matter. We found that vasculature-associated oligodendrocytes (vOLs) are common and are tightly associated with the endothelial basement membrane suggesting that oligodendrocytes should be considered as an integral part of the vasculature microenvironment.

## Materials and Methods

### Animals

Experimental procedures were reviewed and approved by the ethics committee of the University of Bordeaux (CEEA-50) and authorized by the French Ministry of Higher Education, Research and Innovation (agreement number 27726). All animal handling was in agreement with EU directive 2010/63/EU. Mice were purchased from Charles-River or bred in-house. We used male and female C57Bl6/J or CNP-mEGFP mice (Jackson Lab 026105, RRID:IMSR_JAX:026105 (Deng et al., 2014)) that were between 7-12 weeks old. Mice had unlimited access to food and water and were housed in individually ventilated cages on a standard light/dark cycle of 12/12 h. No statistical method was used to pre-determine sample size.

### Cuprizone administration

Male and female mice were fed for 5 weeks with powdered chow alone (control) or powdered chow supplemented with 0.2% cuprizone (bis-(cyclohexanone)-oxal-dihydrazone, C9012, Sigma-Aldrich). Cuprizone-supplemented powder food was replenished every two to three days and supplied in powder food dispensers. After 5 weeks of cuprizone treatment, mice were either sacrificed (average age 12 weeks) or returned to a normal food pellet diet for 7 weeks (average age 22 weeks).

### Tissue preparation for immunohistochemistry

Anesthesia was induced with 3% isoflurane followed by intraperitoneal injection of a ketamine (100 mg/kg)/ xylazine (20 mg/kg) mix. Once mice were deeply anesthetized, we performed transcardial perfusion with PBS supplemented with heparin (12.5 U/ml), followed by freshly prepared 4% paraformaldehyde (Sigma-Aldrich). After fixation, brains were removed and post-fixed between 2 hours to 24 hours in 4% paraformaldehyde. Coronal slices of 50 *μ*m nominal thickness were cut on a vibratome (VT1000S, Leica Microsystems, Germany) or on a freezing microtome (SM2010R, Leica). Cryoprotection of brains was performed by incubating the brains at 4°C in 15% sucrose solution followed by incubation in 30% sucrose solution. Thin sections were stored at 4°C in PBS supplemented with 0.01% NaN_3_ or at −20°C in cryoprotective solution (30% glycerol and 30% ethylene glycol in PBS).

### Immunohistochemistry

For immunolabeling, we selected sections of the primary motor cortex and the primary somatosensory cortex (limb region) based on gross anatomical features. All steps were conducted at room temperature (RT). Free-floating sections were incubated in blocking buffer consisting of 2.5% goat serum, 2.5% bovine serum albumin, and 0.3% triton X-100 in PBS for 1 hour. Primary antibodies (see Table 1) were added to the blocking buffer and sections were incubated overnight. After washing with PBS, sections were subsequently incubated with secondary antibodies for 2 hours in blocking buffer. Sections were washed with PBS and incubated for 5 minutes with 300 nm DAPI (Sigma-Aldrich) and washed afterwards. Tissue sections were mounted on glass slides with Fluoroshield mounting medium (F6182, Sigma-Aldrich) and cover-slipped.

### Confocal Microscopy

Microscopy images were acquired with Leica TCS SP5 confocal microscopes (Leica Microsystems, Germany) controlled by Leica Application Suite software. Tiled z-stack scans covering all cortical layers were acquired either with a 40x 1.30NA objective (HC PL APO CS2 OIL UV, Leica) or a 40x 1.25NA objective (HCX PL APO lambda blue OIL UV, Leica). Higher magnification z-stacks of individual oligodendrocytes located in apposition to blood vessels were taken with a 63x 1.40NA objective (HCX PL APO CS OIL UV, Leica).

### Deconvolution of high-resolution images

High-resolution image stacks were deconvolved using the AutoQuant 3D deconvolution module (version X3.1.2, RRID:SCR_002465, Media Cybernetics, Rockville, MD, USA). We used the adaptive point-spread function method with a theoretical point-spread function and performed 4 iterations with the noise level set to medium.

### Tissue fixation for electron microscopy

For electron microscopy (EM) nine mice were used to optimize the parameters for serial block-face imaging, including perfusion, tissue fixation, tissue preparation, fluorescence conservation, blood-vessel labeling and laser marking. For our final experiments, we performed transcardial perfusion of two mice with PBS (10-15 ml) supplemented with heparin (12.5 U/ml) followed by 25 ml RT fixative solution consisting of 4% PFA and 2.5% glutaraldehyde (both Electron Microscopy Science, Hatfield, PA, USA) in PBS at a flow rate of 2 ml/min. After perfusion the brains were carefully removed from the skull and post-fixed for 24h in the same fixative and then washed several times in PBS before sectioning. Coronal sections of the brains were cut on a VT1000S vibratome (Leica) to a nominal thickness of 100 *μ*m and stored in 0.1 M phosphate buffer (PB). Slices were washed with PB and stored in PB supplemented with 0.02% NaN_3_ until further processing steps.

### 2-photon near-infrared branding for serial block-face imaging

As conventional immunohistochemistry methods do not preserve the ultrastructure, we took advantage of the oligodendrocyte reporter mouse that expresses membrane-tethered EGFP in all oligodendrocyte processes and cell bodies under the CNP promoter. Preliminary experiments confirmed that EGFP fluorescence was preserved after fixation of tissue for EM. Labeling of the vasculature was achieved by incubating 100 *μ*m slices overnight with tomato lectin conjugated with DyLight-649 (Table 1) in PB, which did not require permeabilization. Sections were washed multiple times with PB and transferred to a glass petri dish filled with PB and held with a slice anchor (SHD-27H/15, Warner Instruments, Holliston, MA, USA) for easy recovery of the slices after near-infrared branding. All procedures were performed on a SP5 confocal microscope (Leica) controlled by LAS software (Leica) and equipped with a 25x 0.95NA water-dipping objective (HCX IRAPO, Leica). We performed overview scans at 2048 x 2048 pixels (zoom 1 to 1.5) to identify vOLs and subsequently marked several vOL positions in each slice with laser burns.

For burning laser marks, we used a tunable pulsed 2-photon laser (Mai Tai HP, Spectra Physics/Newport, Irvine, CA, USA) with an average output of ≈1.8 W and wavelength set to 910 nm. 2-photon laser output power (transmission) was not attenuated and kept at 100%, gain was set to 100% and the offset to 73%.

For each identified vOL, the oligodendrocyte cell body was centered under a small horizontal burn mark of ≈20 *μ*m length (XZT mode; digital zoom 30; 512 x 32 pixels/ image; 200 Hz scan speed; 50 lines; average pixel dwell time 10 *μ*s). To identify the position of the laser marks during EM sample preparation, we burned a 300*μ*m long horizontal line (XZT mode; digital zoom 2; 512 x 32 pixels/image; 200 Hz; 1000 lines) and a large square (XYT mode; digital zoom 15; 64 x 64 pixels/image; 50 Hz; 300 frames) above the regions of interest (ROIs). Repetitions were set to match required thickness and line depth (Supplementary Figure 1B,C). Location, width and depth of branding marks were subsequently imaged using a 561 nm laser in normal confocal scan mode, taking advantage of tissue auto-fluorescence on the borders of laser marks.

### Sample preparation for block face electron microscopy

Tissue for serial block-face scanning electron microscopy (SBF-SEM) was prepared similar to previously described (Deerinck et al., 2010). Samples were post-fixed with a 2% paraformaldehyde and 2.5% glutaraldehyde solution (Electron Microscopy Science) in 0.15 M cacodylate buffer (pH 7.4) for 2 hours at RT and subsequently washed five times for 3 minutes in cold 0.15 M cacodylate buffer. To enhance contrast of the tissue, we performed consecutive incubations in heavy metal solutions. Samples were incubated for 1 hour in 2% OsO_4_ containing 1.5% potassium ferrocyanide in 0.15 M cacodylate buffer on ice. After washing five times for 3 minutes in ultrapure water, the samples were incubated for 20 minutes in a freshly prepared thiocarbohydrazide solution (1% w/v in water) at RT. Samples were washed five times for 3 minutes in ultrapure water and then incubated in 2% osmium in water at RT for 30 minutes. After washing 5 times for 3 minutes in ultrapure water the tissue was incubated in 2% uranyl acetate at 4°C overnight. Finally, Walton’s lead aspartate staining was performed for 30 min at 60°C. We prepared a fresh 30 mM L-aspartic acid solution to dissolve lead nitrate (20 mM, pH 5.5), which was subsequently filtered to remove undissolved particles. After final washing steps in ultrapure water, the samples were dehydrated in ice-cold solutions of 30%, 50%, 70%, 90%, and twice in 100% ethanol (anhydrous), and twice in 100% acetone for 10 minutes each. Tissue was embedded in epon by placing the samples in 25% acetone/epon for 2 hours, 50% acetone/epon for 2 hours, 75% acetone/epon for 2 hours and followed by two incubations in 100% epon (overnight, 8 hours). The samples were then transferred to fresh epon resin fand incubated at 60°C for 48 h. Once the resin blocks were hardened, blocks were coarsely cut with a razor blade to generate a pyramidal shaped sample. Embedded samples were mounted on aluminum specimen pins using a silver filled conductive resin (Epotek-Delta microscopies, Mauressac, France). After 24 hours of polymerization at 60°C, the samples were further trimmed with a diamond knife (Diatome, Nidau, Switzerland) on an ultramicrotome (Leica EM UC7). Silver filled conductive resin was used to electrically connect the edges of the tissue to the aluminum pin. The entire sample was sputter-coated with a 5-10 nm layer of gold to enhance conductivity.

### Serial block face scanning electron microscopy (SBF-SEM)

SBF-SEM was performed with a ZEISS Gemini field emission gun SEM300 (Zeiss, Marly-le-Roi, France), equipped with a 3View2XP in situ ultramicrotome (Gatan Inc., Pleasanton, CA, USA). Serial section thickness was set to 30 nm. The block face was imaged with the accelerating voltage set to 1.2 kV and backscattered electrons were detected with an OnPoint Detector (Gatan Inc.) and a pixel dwell time of 6 *μ*s. Images were acquired in high vacuum mode with magnifications and image sizes adjusted to a final pixel size of 10 nm at specimen level.

### Electrophysiology

For electrophysiological investigations of vOLs, we used CNP-mEGFP reporter mice for identification of oligodendrocytes. Mice were deeply anesthetized with a mixture of ketamine (100 mg/kg) and xylazine (20 mg/kg). For some animals, we performed transcardial perfusion with ice-cold preparation solution before decapitation, however, blood vessel visibility was reduced in acute slices. In a subset of experiments, we therefore omitted transcardial perfusion and decapitated deeply anesthetized mice followed by quick removal of the brain. The brain was then submerged in ice-cold preparation solution saturated with carbogen (95% O_2_/5% CO_2_) composed of (in mM) 60 NaCl, 25 NaHCO_3_, 1.25 NaH_2_PO_4_, 2.5 KCl, 100 sucrose, 1 CaCl_2_, 5 MgCl_2_ and 20 glucose. Subsequently, we cut 300 *μ*m para-sagittal slices on a vibratome (VT1200S, Leica, Germany). Slices were collected in carbogen-saturated storage solution composed of (in mM) 125 NaCl, 25 NaHCO_3_, 1.25 NaH_2_PO_4_, 3 KCl, 1 CaCl_2_, 6 MgCl_2_ and 20 glucose. Slices were incubated for 35 minutes at 35°C before being kept in the same solution at RT for the experimental day.

For recordings, slices were transferred to a heated (32 ± 1°C) submerged recording chamber on an upright microscope (Olympus or LN-scope) equipped with infrared illumination and oblique contrast optics for visualization of cells. The recording solution was composed of (in mM) 125 NaCl, 25 NaHCO_3_, 1.25 NaH_2_PO_4_, solution was composed of (in mM) 125 NaCl, 25 NaHCO_3_, 1.25 NaH_2_PO_4_, 3 KCl, 2 CaCl_2_, 1 MgCl_2_ and 25 glucose. Oligodendrocytes were identified in CNP-mEGFP mice by epifluorescence illumination with a 470 nm LED and a GFP filter set (GFP-30LP-B-000, Semrock, Rochester, NY, USA). Vasculature-associated oligodendrocyte cell bodies were identified by switching between epifluorescence and infra-red illumination.

For whole-cell recordings, we used borosilicate glass pipettes (BF150-86-10, Sutter Instrument, Novato, CA, USA) pulled on a vertical puller (PC100, Narishige International Limited, London, UK) to a size of ≈5-7 MΩ. Pipettes were filled with intracellular solution consisting of (in mM) 130 K-Gluconate, 10 KCl, 10 HEPES, 4 Mg-ATP, 0.3 Na_2_-GTP, 10 Na_2_-phosphocreatine with a pH set to 7.25 with KOH and an osmolarity adjusted to 280 mOsm. Satellite oligodendrocytes were recorded with a modified intracellular solution containing 125 mM K-Gluconate and additionally 20 *μ*g/ml glycogen and 0.32 U/*μ*l Ribolock (Thermo-Scientific). Experiments were controlled, and acquired by an integrated patch-clamp amplifier system with SutterPatch software (IPA2, Sutter Instruments, Novato, CA, USA). Recordings were sampled with a minimum frequency of 10 kHz and filtered with a 4-pole Bessel filter set to 5 kHz. After establishing whole-cell configuration in voltage-clamp, we performed recording of membrane potential and membrane characteristics in current-clamp.

### Image analysis software

All image analysis was carried out with FIJI (Schindelin et al., 2012), Digital Micrograph software (Gatan Inc., Pleasanton, CA, USA), Microscopy Image Browser – MIB v.2.702, RRID:SCR_016560 (Belevich et al., 2016), or Imaris version 9.8.0 (Oxford Instruments, Belfast, UK). Imaris was also used to generate 3D reconstruction of images.

### Oligodendrocyte-vasculature distance measurement

The distance separating the oligodendrocyte cell body and blood vessel wall (Figure 1D) was estimated from deconvolved high-resolution images using FIJI. The fluorescence intensity of each channel (CNP and CD31 or Collagen-IV) was measured along a line crossing the contact site (line tool, 1 pixel width) and the distance between the fluorescence peaks was calculated.

**Figure 1.**
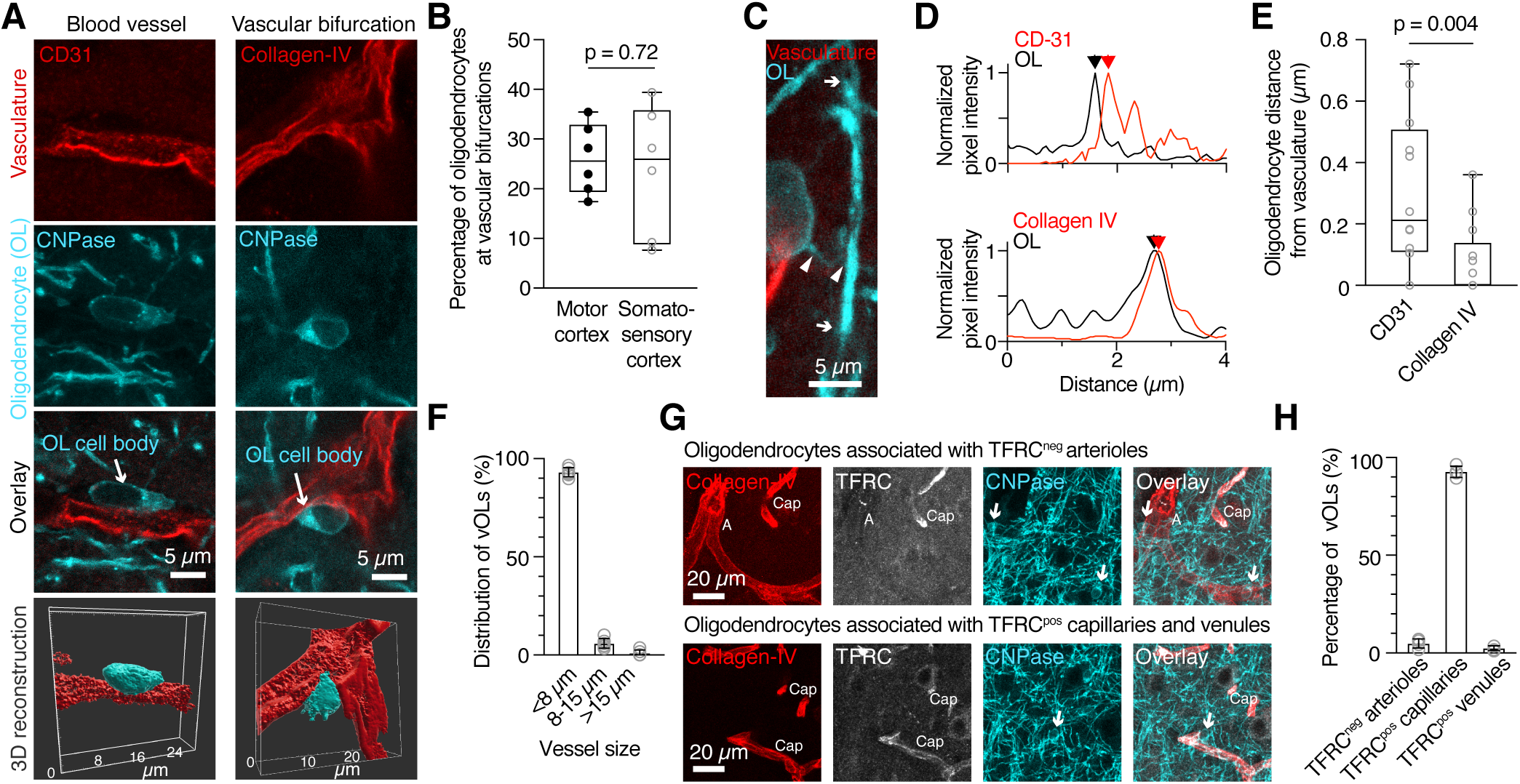
Oligodendrocytes are closely and predominantly associated with capillaries in the cortical gray matter. (A) Representative images of vOLs on blood vessels or vascular bifurcations and corresponding 3D reconstructions. Oligodendrocyte cell bodies (cyan) appear directly associated with the vasculature (red). (B) Quantification of oligodendrocytes on vascular bifurcations from the total population of vOLs (Unpaired t-test, p = 0.72, n = 6 mice/ area). (C) High magnification image of a myelin sheath connected to a vOL. Arrowheads indicate the cell process linking the myelin sheath to the cell body, arrows mark the extension of the internode. The majority of the blood vessel was cut for display. (D) Normalized pixel intensity plots used for quantifying the distance between vasculature and oligodendrocytes. Labeling of either CD31 or collagen-IV shows sub-micrometer distances between blood vessel wall and oligodendrocyte cell body. Maximum peaks of the immunosignal (red and black arrowheads) were used to quantify spacing between immunosignals. (E) Quantification based on (D) reveals a shorter distance for collagen-IV compared to CD31, consistent with blood vessel anatomy (Mann-Whitney test, p = 0.004, n = 12 cells for CD31 and n = 17 cells for collagen-IV from n = 4 mice). (F) Distribution of vOLs at different sized blood vessels shows that the majority is associated with vessels <8 *μ*m, presumably capillaries. (G) Representative images of oligodendrocytes on transferrin receptor negative (arterioles) and positive (capillaries and venules) blood vessels. Arrows point to oligodendrocyte cell bodies. (H) Quantification of blood vessel type reveals that the majority of vOLs are located on transferrin-positive capillaries (n = 6 mice).

### Oligodendrocyte-blood vessel contact area analysis

For accurate reconstructions of oligodendrocyte cell bodies, we segmented the cell outline from deconvolved high-resolution images using MIB software. We exported and rendered the created model as an Imaris surface element and combined the model with the source image in Imaris. The blood vessel surface was reconstructed using the automatic surface creation module of Imaris (settings: smooth; surface detail: 0.120 *μ*m; thresholding: absolute intensity). The surface of contact between the oligodendrocyte soma and blood vessel was created and measured with the surface-surface contact area Imaris XTension (https://imaris.oxinst.com/open/view/surface-surface-contact-area), using the oligodendrocyte surface as primary surface.

### Aquaporin-4 intensity measurements

AQP4 fluorescence was measured from confocal images with line or area ROIs (FIJI) that were placed on AQP4 positive blood vessels, either at the oligodendrocyte-vasculature contact site or outside the contact site.

### Oligodendrocyte number, distribution and density

Cortical organization of oligodendrocytes was analyzed on image z-stacks in FIJI. A grid was overlayed (25 x 25 *μ*m square unit) to the images that was aligned to the surface of the cortex. Oligodendrocytes were manually counted and their coordinates were registered using the point tool and ROI Manager. Cortical layers were determined based on cortical depth and labeling of DAPI and CNP, reflecting cell and myelin densities respectively. An oligodendrocyte was considered as vasculature-associated when the distance separating its cell body from blood vessel wall (measured as described above) was ≤ 1 *μ*m.

### Blood vessel diameter

For each vOL identified in image z-stacks, the diameter of the associated blood vessel was measured with the line tool (1 pixel width, FIJI) within a maximum 20 *μ*m radius around the vOL cell body. Three categories of vessels were defined based on their diameter: <8 *μ*m (cerebral capillaries), 8 to 15 *μ*m (precapillary arterioles/ postcapillary venules) and >15 *μ*m (arterioles/ venules and arteries or veins) (Steinman et al., 2017; Todorov et al., 2020).

### Analysis of myelin and vasculature density

ROIs of 104 *μ*m^2^ ROIs were delimited in each cortical layer from the same images that were used for oligodendrocyte quantification. All analysis was performed with FIJI. Myelin density: A threshold was set for single z-plane images containing myelin labeling (CNPase) and the image subsequently de-speckled to remove 1×1 positive pixels. The total myelin occupying pixels were measured and myelin density was calculated by dividing the myelin positive area by the total area of the ROI.

Vasculature density: We performed maximum intensity z-projections of images with collagen-IV labeling. An image threshold was set and positive outliers (<3 pixels) resulting from image acquisition noise were removed. Vasculature density was calculated by dividing the total blood vessel area by the volume of the ROI.

### Image processing of electron microscopy images

Automatic image registration and alignment of the raw image stack was performed with Digital Micrograph using the forward running image as reference and a subpixel alignment accuracy to remove drift from image acquisition. The resulting stack was cropped to an area common for all images. All further image processing was performed on tif images. We used MIB for manual segmentation of the blood vessel lumen, basement membrane, oligodendrocyte cell body and main protruding processes using a drawing tablet and pen (Cintiq pro 5, Wacom, Saitama, Japan) as input device. We created independent models of each element from which we generated 3D models in Imaris (9.8.0).

### Electrophysiological analysis

Data were exported from Sutter Patch and analyzed offline using pClamp 10.1 (Molecular Devices LLC, San Jose, CA, USA). The resting membrane potential (V_m_) was determined from baseline segments of current-clamp recording traces with zero-current injection. Input resistance (R_in_) was estimated by fitting the linear range of voltage responses to 400 pA step current injections ranging from −800 to 400 pA. Resting conductance (Gm) was calculated as *G_m_* = ΔI/ΔV where ΔV was based on a 10 mV hyperpolarizing voltage step from holding potential. For I-V relationship analysis, we measured the steady-state current (indicated in Figure 4C) as a response to voltage steps between −10 mV and −130 mV. Principal component analysis was performed by using the R (R Core Team, 2021) prcomp function with standard parameters and inclusion of four electrophysiological variables: resting membrane potential, conductance, input resistance and capacitance. All reported voltage values were corrected for a liquid junction potential of −15 mV as previously determined (Battefeld et al., 2016).

### Statistical analysis

Data were statistically analyzed in Prism 9 (Version 9.3, GraphPad Software, San Diego, CA, USA). Data were tested for normal distribution with a Shapiro-Wilk test. For comparison between two groups, non-normally distributed data were analyzed by a Mann-Whitney test and normally distributed data were tested with either a paired or unpaired t-test. Comparisons of more than two groups were performed using a non-parametric Kruskal-Wallis test. A probability of p < 0.05 was considered statistically significant. In the text, data are given as mean ± standard error of the mean.

### Data availability statement

The data that support the findings of this study are available from the corresponding author upon reasonable request.

## Results

### Oligodendrocytes associate with the vasculature in the neocortex

Early work noted the presence of oligodendrocytes close to blood vessels in the cortex (Cammermeyer, 1960; del Rio Hortega, 1928), but occurrence and precise location on the vascular network remained largely unexplored. Using immunohistochemistry, we determined the location of mature oligodendrocytes on the brain vasculature labelled by the endothelial cell marker CD31 or the basement membrane component collagen-IV in 8-week-old mice (Figure 1A). Oligodendrocyte cell bodies contacted blood vessels, either along unbranched sections or at vascular bifurcations (Figure 1A). Additionally, 22% of vOLs (309 out of 1392 vOLs, n = 6 mice) were located at vascular bifurcations in both somatosensory and motor areas (Figure 1B). Next, we traced processes from the oligodendrocyte cell body to myelin sheaths, confirming that vOLs are myelinating (n = 4 mice, Figure 1C).

Since vOLs are in close contact with the vasculature, we measured the distance between oligodendrocyte cell bodies and the vessel wall, based on fluorescence intensity peaks of their respective markers (Figure 1D). In line with the blood vessel anatomy, vOL cell bodies were further away from the endothelial cell marker CD-31 (300 ± 247 *μ*m, n = 12) compared to the basement membrane marker collagen-IV (80 ± 127 *μ*m, n = 17, Mann-Whitney test p = 0.004, Figure 1E). These short distances indicated a possible direct contact with the vasculature.

The vasculature is divided into arterial, capillary and venous networks, which differ in terms of size, structure and function. We asked whether vOLs preferentially locate to specific parts of the vascular network. Based on the blood vessel diameter at the contact site, the majority (93% ± 2.4%) of oligodendrocytes was located on vessels with a diameter <8 *μ*m (969 of 1049 cells, n = 6 mice, Figure 1F) corresponding to capillaries (see methods for definition). Only a few oligodendrocytes were located on larger diameter vessels (>8 *μ*m 66 of 1049 cells; >15 *μ*m 14 of 1049 cells). These results are in line with volumetric data from mouse cortex, showing that the majority of vessels are micro-vessels (Todorov et al., 2020). To determine if vOLs on larger vessels are found on the arterial or venous part of the vascular network, we next labelled blood vessels with transferrin receptor (Tfrc), which is expressed in capillaries and veins, but not in arteries (Vanlandewijck et al., 2018). Similar to the size distribution analysis, the majority of oligodendrocytes were located on Tfrc-positive capillaries. In addition, few oligodendrocytes were found on either Tfrc-positive venules or Tfrc-negative arterioles (Figure 1G, H). In summary, our data establish that in the neocortex, vOL cell bodies are in close contact with different blood vessel types and mostly associate with capillaries.

### Oligodendrocytes directly contact the vascular basement membrane

Although our immunohistochemistry experiments revealed that oligodendrocytes are in close contact with the vasculature, the measured distance between oligodendrocytes and blood vessels was at or below the resolution limit of standard optical microscopy. We therefore investigated this contact further with EM. To specifically target vOLs, we used a correlative light and EM approach in which we marked the tissue location of vOLs with 2-photon near-infrared branding (Figure 2A and Supplementary figure 1A-G) ahead of sample preparation for EM (Bishop et al., 2011). Using SBF-SEM of laser marked vOLs, we established that the oligodendrocyte cell body is in direct contact with the basement membrane throughout several z-sections (n = 2 cells from 2 mice, Figure 2B). Processes of the imaged vOL were contacting myelin sheaths in adjacent serial sections (Figure 2C), confirming that vOLs myelinate axons. In line with previous observation (Peters et al., 1991; Shapson-Coe et al., 2021), we observed that the cytoplasm of the oligodendrocyte cell body was darker than surrounding tissue elements (Supplementary figure 1H). A 3-dimensional reconstruction (Figure 2D) showed that the oligodendrocyte soma and the vascular basement membrane are in direct contact over a large area, surrounded with what is likely to be astrocytic endfeet. To estimate the contact area, we used confocal images of vOLs and blood vessels. The average contact area was 36.5 ± 23.1 *μ*m^2^, representing about 13.6 ± 6.7% (n = 13 cells, n = 3 mice) of the total oligodendrocyte surface area (Figure 2E,F).

**Figure 2.**
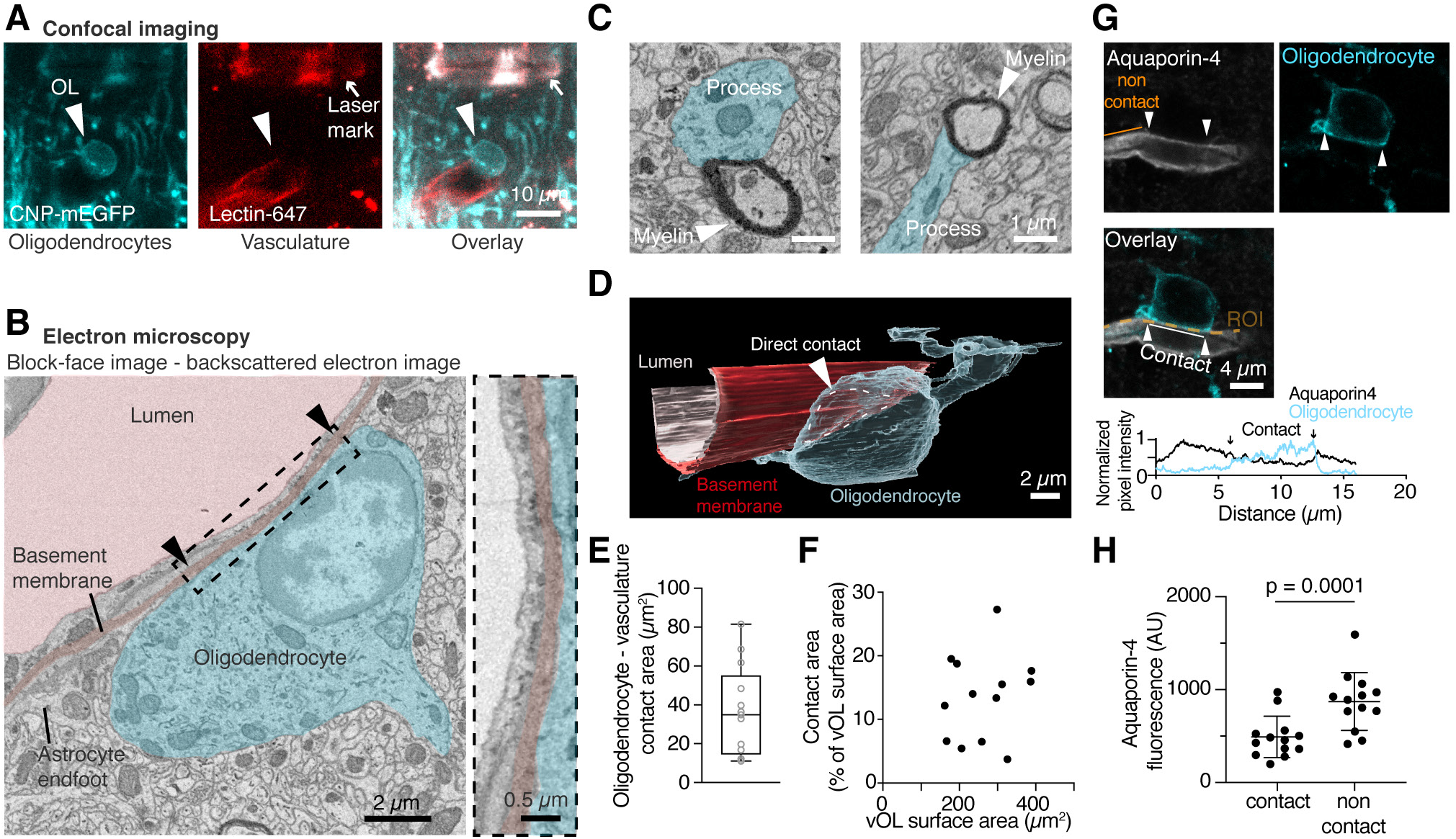
Oligodendrocytes are in direct contact with the vascular basement membrane. (A) Confocal images of a vOL (cyan, arrowhead) on a blood vessel (red). The location was marked with 2-photon branding and the created laser mark is well visible using green excitation light (see arrow). (B) Single-plane EM image of the same oligodendrocyte as in (A) showing the direct contact between the cell body (light cyan,) and the basement membrane (dark red). Arrowheads denote the length of the contact and the boxed area is displayed at higher magnification. (C) Two example EM images showing traced processes (light cyan) ending in compact myelin. (D) 3D rendering of the reconstructed vOL, the basement membrane and the blood vessel lumen. The oligodendrocyte cell body closely follows the blood vessel shape. (E) Quantification of oligodendrocyte-vasculature contact area from 3D confocal image reconstructions. (F) Contact area with the vasculature is not correlated with the oligodendrocyte total surface area (Pearson r^2^ = 0.036, p = 0.53, n = 13). (G) Single plane confocal images showing aquaporin-4 labeling of the vasculature (grey) and oligodendrocyte cell body (cyan) labeling in a CNP-mGFP mouse. Bottom: Fluorescence intensity plots for both channels, showing a reduction of Aquaporin-4 labeling intensity at the contact site. (H) Quantification of the aquaporin-4 fluorescence and comparison between contact and non-contact sites show a reduced intensity of aquaporin-4 signal at the contact sites (Paired t-test, p = 0.0001, n = 13 cells from 4 animals).

As we could not unequivocally identify astrocytic endfeet within the volumetric EM images, we specifically labelled endfeet with aquaporin-4 (AQP4, Figure 2G) and assessed the presence of AQP4 labeling at or around the contact site with confocal microscopy. We observed a lower fluorescence intensity of AQP4 at the contact site, compared to the blood vessel segments that were not in contact with the oligodendrocyte (paired t-test, p = 0.0001, n = 13, Figure 2G,H). This suggests that astrocytic endfeet are contacting vOL cell bodies, but are reduced in size or not present directly at the contact site in line with our electron microscopy data. We conclude that vOLs are in immediate proximity to astrocytic endfeet that surround the vasculature and that vOLs establish a direct contact with the vascular basement membrane.

### Distribution of vasculature-associated oligodendrocytes in the neocortex

Since oligodendrocytes are non-uniformly distributed across the cortex (Tomassy et al., 2014), we asked whether vOLs follow a specific pattern of distribution across cortical layers. To map the vOL population, we labelled the vasculature with CD31 or collagen-IV and oligodendrocytes and myelin with CNPase (Figure 3A). Vascular density appeared uniformly distributed (Figure 3B), whereas oligodendrocytes and myelin density increased with cortical depth, in line with previous work (Tomassy et al., 2014). The increase in myelin density was positively correlated with an increase in oligodendrocyte cell body density (Figure 3C), thus oligodendrocyte cell body location and density can be used to infer myelination density.

**Figure 3.**
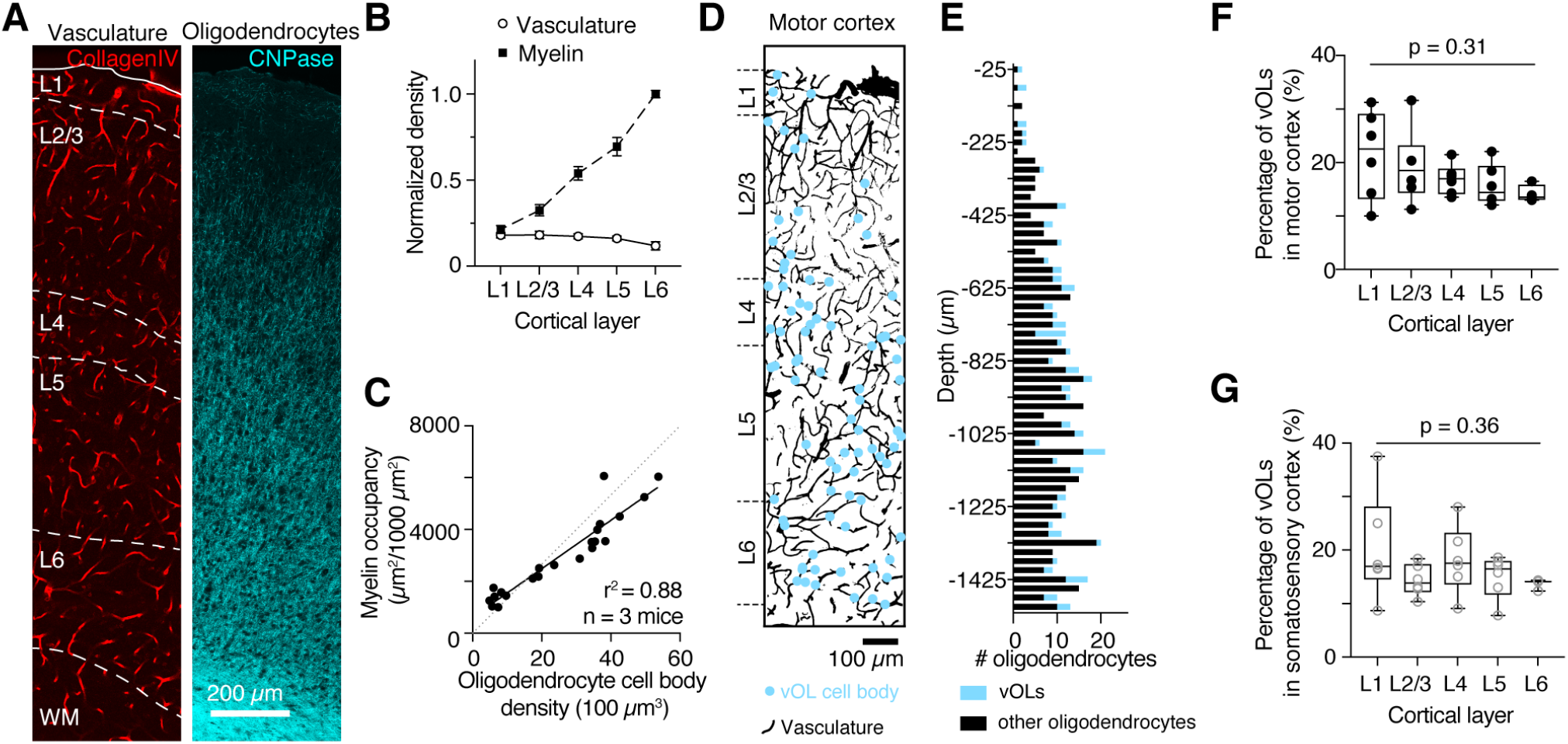
Oligodendrocytes associate with the vasculature in all cortical layers. (A) Representative single plane confocal images of motor cortex showing collagen-IV labelled vasculature (red) and CNPase labelled oligodendrocytes (cyan). (B) Quantifications reveal that vascular density remains constant throughout the cortex, whereas myelin density increases towards deeper layers. Data were normalized to myelin density in layer 6 (n = 3 mice). (C) Oligodendrocyte cell body density is highly correlated with myelin immunosignal (myelin occupancy), confirming that oligodendrocyte cell body location and density can be used to infer myelination density. Data from n = 3 mice were fit with a linear function. (D) Vasculature (black) from a z-projected cortex volume (same as in A) overlayed with vOL locations (cyan). (E) Plot showing oligodendrocytes occurrence within the volume in D (25 *μ*m bins). Vasculature-associated oligodendrocytes are highlighted in cyan. (F, G) Throughout all cortical layers, the proportion of vOLs remains constant in the motor cortex (F) and somatosensory cortex (G). Data was tested with a Kruskal-Wallis test (n = 6 mice for layers 1 to 5, and n = 3 mice for layer 6).

Next, we determined vOL locations in all cortical layers (Figure 3D). Similarly, to the overall increase in oligodendrocyte density in deeper cortical layers (Supplementary Figure 2B and 2D), vOLs increased in occurrence with cortical depth (Figure 3E and Supplementary Figure 2C and 2E). As a result, the proportion of vOLs remained stable across all neocortical layers. In the motor cortex, vOLs represented 17.7 ± 1.1% of oligodendrocytes (n = 6 mice, Kruskal-Wallis test p = 0.31, Figure 3F) and 16.6 ± 1.2% in the somatosensory cortex (n = 6 mice, Kruskal-Wallis test p = 0.36, Figure 3G) across all cortical layers. The total percentage of vOLs across the whole cortex was similar in primary motor and somatosensory areas (Mann-Whitney test, p = 0.5, n = 6 mice). The increase of vOLs with cortical depth was not correlated with the vascular density (Pearson correlation, R^2^ = 0.21, p = 0.1, n = 3 mice). As brain regions differ in their myelination patterns, we examined the presence of vOLs in other brain areas. Analysis of the hippocampal CA1 region and the cerebellar anterior lobe confirmed the presence of vOLs in both regions (Supplementary Figure 2F, G). Quantifications showed that vOLs represent 16.6 ± 2.3% of the oligodendrocyte population in the hippocampus (n = 3 mice) and 20.8 ± 2.8% in the cerebellum (n = 4 mice, Supplementary Figure 2H). In summary, one out of six oligodendrocytes associates with the vasculature across different brain regions. We conclude from these experiments that vOLs are frequent, and an important component of the vasculature micro-environment in the adult brain.

### Vasculature-associated oligodendrocytes are regenerated during remyelination

It was previously reported that OPCs associate with the vasculature during remyelination (Niu et al., 2019) and differentiate to mature oligodendrocytes to compensate for oligodendrocyte loss in the lesion. However, during remyelination newly formed oligodendrocytes appear in different locations and display a different myelination pattern (Bacmeister et al., 2020; Orthmann-Murphy et al., 2020; Snaidero et al., 2020). To assess potential changes of the vOL population during remyelination, mice were fed with cupri-zone-supplemented food for 5 weeks to induce demyelination and subsequently returned to normal food for 7 weeks (Figure 4A). We first analyzed the extent of cortical demyelination in the motor cortex after 5 weeks of demyelination (Figure 4B, C). Compared to control mice, the cortex of cuprizone-fed mice was almost completely devoid of oligodendrocytes except for some oligodendrocytes in layer 2/3 and a few oligodendrocytes close to the white matter. Quantifying the density of the remaining oligodendrocytes in layer 2/3 showed that after 5 weeks of demyelination, only 8 ± 1 oligodendrocytes per 100 *μ*m^3^ remained compared to 27 ± 5 oligodendrocytes per 100 *μ*m^3^ in control cortex (Figure 4C, unpaired t-test, p = 0.023). Even though we could not distinguish between new oligodendrocytes formed during demyelination or oligodendrocytes that survived the cuprizone treatment, these results confirmed that cuprizone treatment was highly effective in reducing cortical oligodendrocytes.

**Figure 4.**
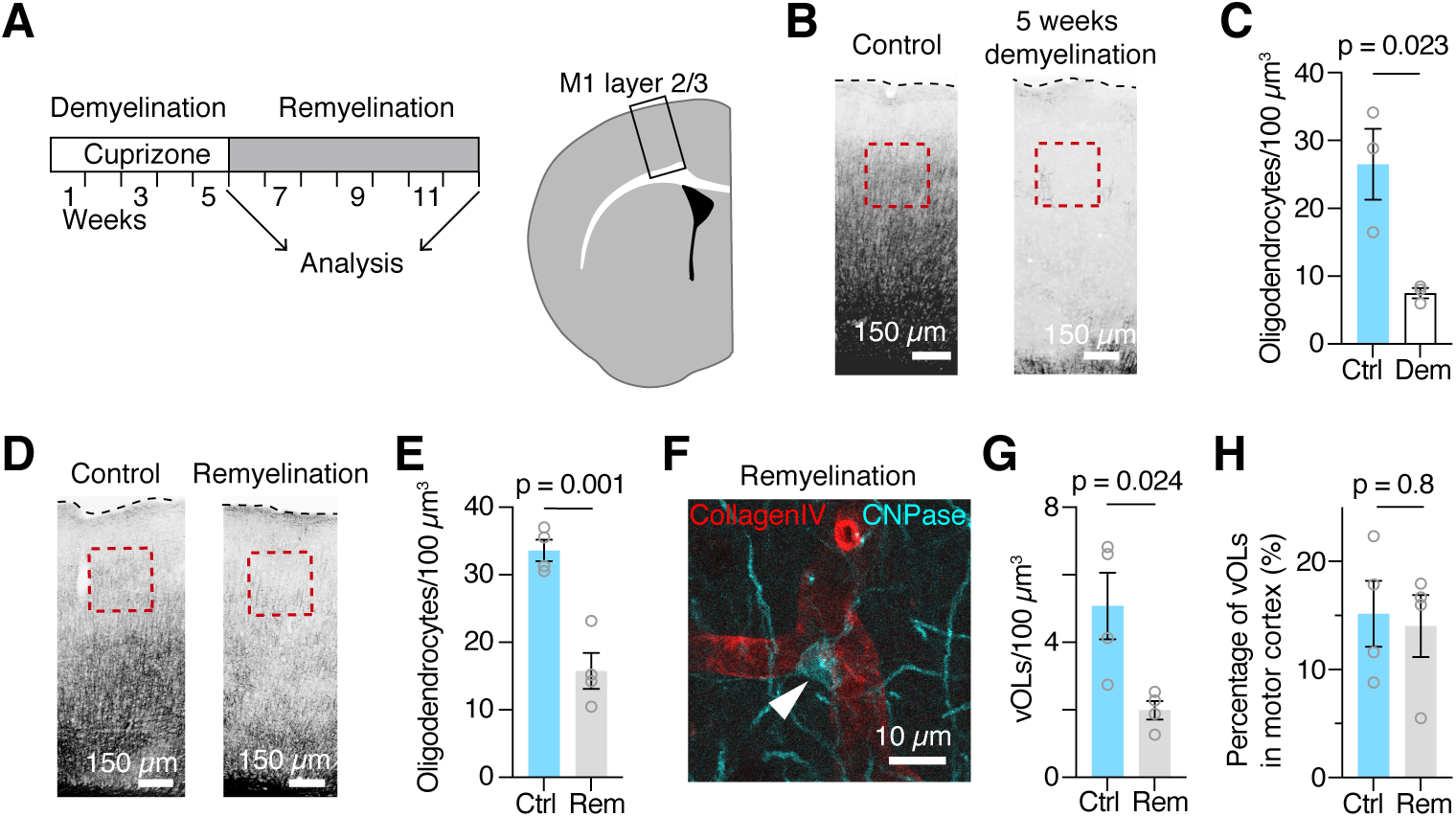
Remyelination re-establishes vascular association of oligodendrocytes. (A) Left: Schematic of the experiment with indicated time points of analysis after 5 weeks of demyelination and 7 weeks of remyelination. Right: Schematic overview and locations of the microscopy images shown in B and D (black box). (B) 5 weeks of cuprizone treatment leads to near complete demyelination of the motor cortex in comparison to control cortex. Red boxes indicate the zone of quantification in layer 2/3, a dotted line denotes the cortex surface. (C) Quantifications of the oligodendrocyte density after 5 weeks of demyelination in layer 2/3, compared to age matched controls. Animals were 12 weeks old at time of analysis. (D) Example overview images of the cortex from age matched control and remyelinated mice. In the remyelinated cortex, myelin still appears less homogeneous compared to control. Red boxes indicate the area of quantification in layer 2/3. (E) Quantifications in layer 2/3 show that after 7 weeks of remyelination, total oligodendrocyte density is still reduced in motor cortex compared to age matched control mice. Single data points represent averaged data from one animal. Age of mice at time of analysis is 22 weeks. Unpaired t-test, p = 0.001. (F) Example of a vOL in motor cortex (arrow head) after 7 weeks of remyelination. (G) Quantifications of vOL density during remyelination show a reduction compared to control. Single data points represent data from one animal. Unpaired t-test, p = 0.024, n = 4 animals for control and for remyelination. (H) Quantifications reveal that the percentage of vOLs during remyelination is comparable to the control motor cortex. Unpaired t-test, p = 0.8, n = 4 animals for control and for remyelination.

To investigate whether mature oligodendrocytes associate with the vasculature during remyelination, we analyzed the layer 2/3 of the motor cortex after 7 weeks of remyelination (Figure 4D). We found that oligodendrocytes in these mice were still reduced by 50 ± 9% (n = 4 mice) compared to age matched control mice (n = 4 mice, Figure 4E, unpaired t-test, p = 0.001). In the remyelinated cortex vOLs were present (Figure 4F), but reduced by 63 ± 7% compared to age matched control mice (n = 4 mice in control and remyelination, Figure 4G). While the overall number of vOLs was decreased, the proportion of vOLs in the remyelinated motor cortex was comparable to the control cortex (Figure 4H).

In the remyelinating motor cortex, vOLs occur in the same proportion as in control mice, suggesting a homeostatic regulation of this oligodendrocyte population.

### Physiological properties of vasculature-associated oligodendrocytes

Given the role of oligodendrocytes in potassium buffering (Battefeld et al., 2016; Larson et al., 2018) and the effect of extracellular potassium in vasodilation (Horiuchi et al., 2002; Longden et al., 2017) we explored intrinsic membrane properties of vOLs and compared them to satellite oligodendrocytes (Battefeld et al., 2016) that are attached to neuronal somas. We performed whole-cell recordings from identified vOLs and satellite oligodendrocytes in acute slices of the neocortex (Figure 5A, B). We determined the resting membrane potential of vOLs (−86.5 ± 3.8 mV, n = 22 cells) from current-clamp recordings (Figure 5C), which was comparable to satellite oligodendrocytes (−84.4 ± 3.2 mV, n = 17 cells, unpaired t-test p = 0.08, Figure 5D). Input resistance was also similar between the two oligodendrocyte subtypes (Mann-Whitney test, p = 0.66, Figure 5E).

**Figure 5.**
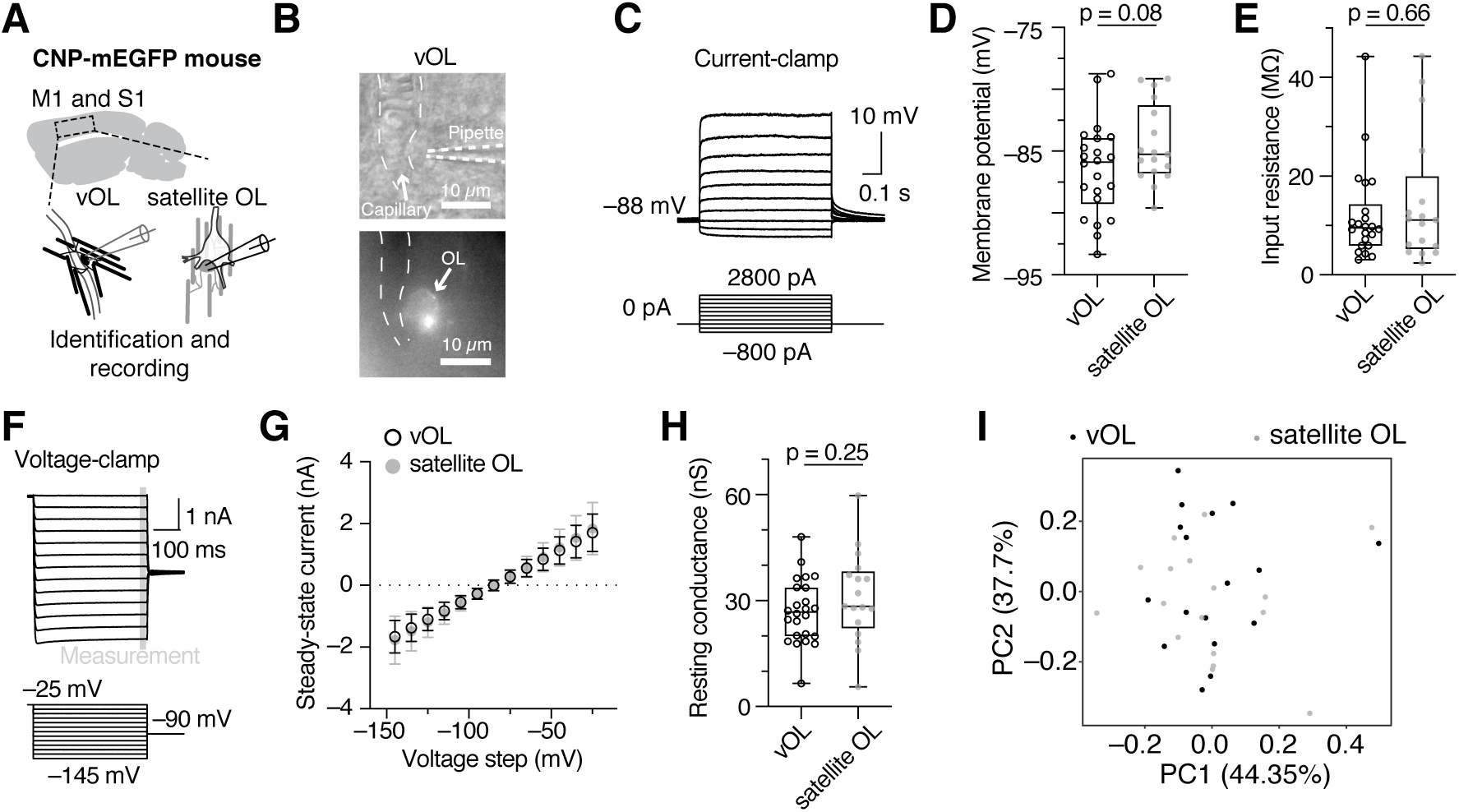
Physiology of vOLs is similar to gray matter satellite oligodendrocytes. ((A) Schematic of the recording location in somatosensory and motor cortex. (B) Top: Example image of a vOL with patch-clamp pipette in oblique contrast. Bottom: The corresponding GFP fluorescence image used to identify the oligodendrocyte. (C) Representative example traces from current-clamp whole-cell recordings of a vOL from which the resting membrane potential and input resistance were determined (D) Resting membrane potential does not differ between vOLs and satellite oligodendrocytes (unpaired t-test, p = 0.36, n = 12 and n = 21 cells). (E) The input resistance was not different between vOLs and satellite oligodendrocytes (Mann-Whitney test, p = 0.65, n = 17 and n = 22 cells). (F) Example whole-cell voltage clamp recording of a vOL with corresponding command voltages. The steady-state measurement is indicated and displayed in G. (G) Current-voltage (I-V) relationship of vOLs compared to satellite oligodendrocytes shows a linear I-V relationship for oligodendrocytes from both locations in the steady-state. (H) Resting conductance calculated from membrane potential is similar between vOLs and satellite oligodendrocytes (unpaired t-test, p = 0.25, n = 17 and n = 24 cells). (I) Principal component analysis based on the electrophysiological parameters shows no separation of the two groups of oligodendrocytes.

We next recorded in voltage-clamp to determine current-voltage relationship over a range of voltage commands. Both oligodendrocyte subtypes had a linear response to command voltages (Figure 5F and 5G) and vOLs had a high resting conductance of 30.9 ± 3.1 nS (n = 24 cells) similar to satellite oligodendrocytes (unpaired t-test p = 0.25, Figure 5H). In line with these data, principal component analysis of the electrophysiological parameters showed no separation of the two anatomically distinct oligodendrocyte populations (Figure 5I).

In summary, vOLs have a similar electrophysiological profile compared to satellite oligodendrocytes in the gray matter, suggesting that these anatomically distinct subpopulations are not clearly identifiable by their electrophysiological properties.

## Discussion

In this study we show that mature oligodendrocytes frequently and closely associate with the vasculature. As the position of oligodendrocyte cell bodies is very stable during lifetime (Hill et al, 2018), we suggest that vOLs are a component of the brain vasculature microenvironment. Given their intermediate location between blood vessels, axons and other oligodendrocytes, vOLs might play essential roles in the central nervous system function.

### Mature oligodendrocyte association with the vasculature

Although the association of oligodendrocytes with blood vessels has been described almost a century ago (del Rio Hortega, 1928) this observation was only explored or noted in few further studies (Cammermeyer, 1960; Vanlandewijck et al., 2018; Vinet et al., 2010). Here, we present evidence based on (i) immunohistochemistry, and (ii) correlative light and volume EM, that vOLs are common and establish a direct contact with the basement membrane of the vasculature (Figure 1 and 2).

It should be noted that chemical fixation and dehydration result in tissue shrinkage and a reduction of the extracellular space around the vasculature (Korogod et al., 2015) suggesting that the cellular arrangement could be more loose (Figures 1 to Figure 4). However, we consistently found vOLs in living tissue in electrophysiology experiments (Figure 5). Our data further support the recent proposition that some oligodendrocytes are physically attached to the vasculature, concluded from a single-cell sequencing data set obtained from purified vessel-associated cells (Vanlandewijck et al., 2018). Moreover, we show that vOLs represent a significant and constant proportion of oligodendrocytes in different brain regions, expanding recent findings of OPC interaction with the vasculature (Chen et al., 2021; Niu et al., 2019; Tsai et al., 2016; Yuen et al., 2014) to mature oligodendrocytes.

### Wnt-dependent association of oligodendrocytes with the vasculature

To date, the Wnt signaling pathway is one of the best identified signaling mechanisms that is involved in vasculature-OPC interaction. It has been demonstrated that physical OPC-vasculature interaction relies on Wnt-dependent signaling via the activation of Cxcr4 receptor in OPCs. Inactivation of this pathway, leads to detachment of OPCs from blood vessels and subsequent OPC differentiation (Tsai et al., 2016). This model is further supported by the recent discovery of OPC perivascular clusters in multiple sclerosis tissue, which has been linked to Wnt pathway overactivation in OPCs (Niu et al., 2019).

The existence of vOLs is in contrast with these earlier works, as vOLs might be generated from non-detached OPCs. Notably, vOLs were regenerated during remyelination (Figure 4), further supporting the possibility that vOLs could originate from OPCs that remain on blood vessels. This reasoning could indicate functional heterogeneity in response to Wnt signaling and its various downstream signaling pathways within the oligodendrocyte lineage (Niehrs, 2012). Local increases in Wnt may alternatively originate from other sources as astrocytes that release Wnt to maintain endfeet integrity (Guérit et al., 2021). As a consequence, a sustained Wnt tone from astrocytes might inhibit the detachment of some OPCs. However, as we observe one out of six oligodendrocytes at the vasculature, the proposed model of OPC detachment (Tsai et al., 2016) remains plausible for the majority of OPCs.

Instead, association of oligodendrocytes with the vasculature could also be Wnt-independent, secondary to detachment, or regulated through other factors. For example oligodendrocytes could be intrinsically primed to attach to the basement membrane via laminin-α2 (Lama2), which is expressed in oligodendrocytes and plays a role in myelination and OPC maturation (Marques et al., 2016; Relucio et al., 2012; Vanlandewijck et al., 2018). Moreover, laminin-α2 has been shown to be required for astrocytic endfeet maintenance and attachment to the basement membrane (Menezes et al., 2014) and might therefore play a similar role in vOLs, but this remains to be further investigated.

### Potential functions of vasculature-associated oligodendrocytes

Oligodendrocytes are heterogeneous (reviewed in Seeker and Williams, 2021) and can be classified based on their developmental origin (Kessaris et al., 2006), transcriptomic status (Floriddia et al., 2020; Marques et al., 2016), anatomical organization (del Rio Hortega, 1928; Floriddia et al., 2020; Peters et al., 1991; Takasaki et al., 2010) or functional impact (Battefeld et al., 2016). Whether the population of vOLs has distinct functional properties remains to be addressed in future functional studies. In the following paragraphs we consider how the direct association of vOLs with the vasculature could influence oligodendrocyte, blood vessel and ultimately neuronal functions.

First, oligodendrocytes provide metabolites, such as lactate and glucose, to neurons to maintain ion gradients during axonal action potential firing (Attwell and Laughlin, 2001; Fünfschilling et al., 2012; Meyer et al., 2018; Saab et al., 2016). In this respect, vOLs are well positioned to directly import glucose from endothelial cells and respond rapidly to increased metabolite demand without an intermediate transfer from astrocytes. This metabolic coupling could be particularly relevant in learning conditions in which increased energy demand is revealed by functional hypoxia (Butt et al., 2021).

Second, oligodendrocytes participate in potassium buffering (Battefeld et al., 2016; Larson et al., 2018) and given the role of extracellular potassium in pericyte relaxation and subsequent dilation of capillaries (Horiuchi et al., 2002; Li and Puro, 2001), vOLs could modulate vascular tone by locally regulating extracellular potassium. Vasculature-associated oligodendrocytes are mainly found on capillaries (Figure 1) and potassium increase around capillaries has been shown to induce upstream dilation of arterioles *in vivo* (Longden et al., 2017). Notably, we observed that 22% of vOLs are located at vasculature bifurcations (Figure 1), which could reflect a role in potassium-dependent vasodilation of higher order blood vessels. Moreover, capillary dilation has also been observed in response to a physiological stimulus or glutamate application (Franco et al., 2022; Hall et al., 2014). Whether vOLs interfere with capillary dilation by modulating extracellular glutamate via glutamate transporter activity (Hamilton et al., 2016; Kukley et al., 2010) remains to be explored.

Third, the existence of a direct contact between vOLs and blood vessels allows a quick response to vasculature derived signals, e.g., the release of endothelin by endothelial cells, which regulates myelin sheath numbers in response to neuronal activity (Swire et al., 2019). However, oligodendrocytes might respond to endothelin only during short periods of maturation and myelin formation (Swire et al., 2019).

Fourth, oligodendrocytes are the brain cells with the highest amount of iron, which is necessary for myelination (Benkovic and Connor, 1993). The majority of vOLs were located on transferrin receptor positive blood vessels (Figure 1G,H) and vessel-associated oligodendrocytes have been found to highly express transferrin, which is required for cellular iron import in the brain (Vanlandewijck et al., 2018). As a result, vOLs may act as an entry point for iron into the gap-junction-coupled oligodendrocyte network.

### Implications in pathologies

Vasculature changes, including alterations of the blood-brain barrier, have been implicated in several neurodegenerative diseases (reviewed in Sweeney et al., 2018). Although vascular dysfunctions have been noted in multiple sclerosis, their relation and impact on disease progression are unclear (D’haeseleer et al., 2011). Notably, type 2 gray matter lesions often occur around the vasculature (Geurts et al., 2009), where early signs of blood-brain barrier disruption are also found (Cramer et al., 2015). In the experimental autoimmune encephalomyelitis model, oligodendrocytes have been identified to present antigens (Falcão et al., 2018) and altered oligodendrocyte heterogeneity has been found in multiple sclerosis (Falcão et al., 2018; Jäkel et al., 2019). It remains to be investigated whether vOLs could be more vulnerable than other oligodendrocytes, as vOLs could either be an early target during immune cell infiltration into the brain parenchyma (Lopes Pinheiro et al., 2016) or play a role in initiating immune cell contact as antigen-presenting cells (Falcão et al., 2018).

Oligodendrocyte-vasculature interaction has also been shown in white matter injury, where secretion of metalloproteinase-9 by oligodendrocytes promotes angiogenesis (Pham et al., 2012), a mechanism that would be especially relevant for vOLs. In conclusion, vOLs might contribute to oligodendrocyte vulnerability and vasculature changes in pathological conditions.

### Outlook and conclusion

We focused on identifying the fundamental characteristics of vasculature interaction with mature oligodendrocytes. Astrocyte interaction with the vasculature is well established, and recent studies show that microglia and OPCs also interact with blood vessels (Bisht et al., 2021; Mondo et al., 2020; Tsai et al., 2016; Yuen et al., 2014). Here, we provide evidence that mature oligodendrocytes interact with blood vessels as well, suggesting a general crosstalk of all glial cells with the vasculature.

Our study presents the basis for future studies of vasculature-oligodendrocyte interaction in both physiology and pathology. Vasculature alterations and the susceptibility of oligodendrocytes in several neurodegenerative diseases show the importance of further addressing this intricate cellular interaction.

## Acknowledgments

This study was funded by a Bordeaux Neurocampus Startup grant provided by the Region Aquitaine (2018.599), IDEX ATTRACTIVITE Chaires Neurocampus of the University of Bordeaux and a grant from the French National Research Agency (ANR-21-CE16-0019-01) to AB. This study received financial support from the French government in the framework of the University of Bordeaux’s IdEx “Investments for the Future” program GPR BRAIN_2030. Imaging was performed at the Bordeaux Imaging Center a member of the FranceBioImaging national infrastructure (ANR-10-INBS-04). We also thank the PIV-EXPE of the University of Bordeaux. We thank Sharlen Moore and Jean-Luc Morel for valuable discussions.

## Author contributions

Conceptualization: AB with input from JP; Investigation: JP, MB, EG, AB; Formal analysis: JP, MB, FSRT, ML, AB; Writing Original Draft: JP, AB; Writing – Review& editing: JP and AB with input from all authors; Visualization JP, AB; Supervision EG, AB; Funding acquisition: AB. All authors approved the final manuscript.

## Notes

### Competing Interest Statement

The authors have declared no competing interest.

