## Supplemental Figures for "A population of gray matter oligodendrocytes directly associates with the vasculature"

**This file includes:**

Figures S1 and S2

Supplementary Figure 1

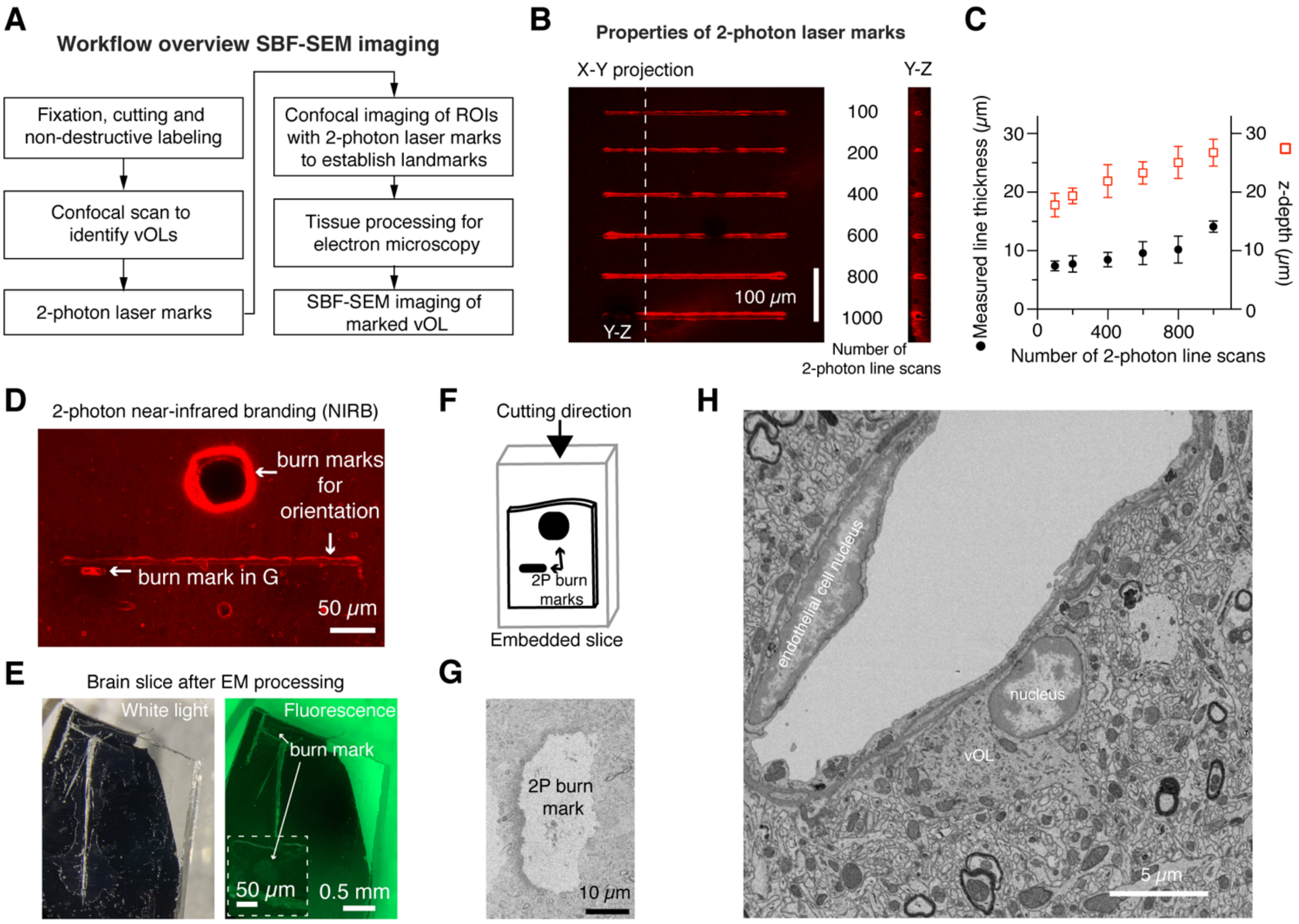


**Supplementary Figure 1: SBF-SEM imaging workflow and experimental steps** (A) Workflow of near-infrared branding guided serial block-face imaging. (B) 2-photon tissue burn lines for estimating the laser power needed to achieve good tissue marking for EM ROI re-localization. (C) Quantification of the laser marks in thickness and depth, used as reference for subsequent 2-photon laser burns. (D) Example image of laser marks used for final marking. A large squared burn mark was made to identify the ROI location under low magnification. (E) Tissue block after the tissue was treated for EM. In ambient light the tissue block is black, prohibiting identification of burn marks. Fluorescent illumination (right) allows identification of the burn mark under binocular observation. The inset shows the burn mark at higher magnification. (F) Schematic of the mounted tissue on the imaging pin. The cutting direction for serial sectioning is indicated. (G) Example of a low resolution backscattered EM image of a small 2-photon branding mark. Small branding marks were used to identify the start and location of the ROI in EM images. (H) Full field of view of the image shown in Figure 2B without pseudo-coloring. The darker cytoplasm of the oligodendrocyte is visible in this representation.

Supplementary Figure 2

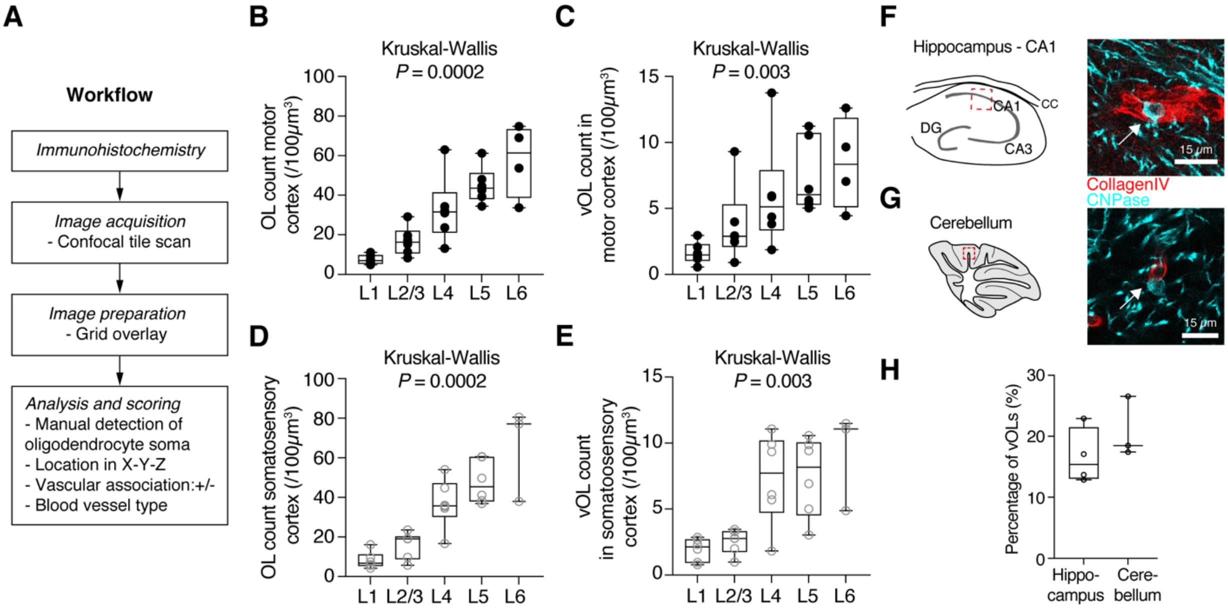


**Supplementary Figure 2: Analysis workflow and additional analysis of vOLs**

(A) Workflow of immunohistochemistry experiments and subsequent analysis steps to determine cortical oligodendrocyte distribution. (B) Total oligodendrocyte density in each layer of the motor cortex (MC) of 9-week-old mice. Each datapoint represent averaged data from one mouse. Data for CD31 and collagen-IV labelling experiments were summarized. (C) Box plot showing density of vOLs in all cortical layers of the motor cortex. (D) Oligodendrocyte density in the somatosensory cortex of 9-week-old mice. Data points contain vasculature labelling with CD31 and collagen-IV. (E) Quantifications of vOL density in the somatosensory cortex for the different layers. (F) *Left:* Schematic of analysis location in the hippocampal CA1 area. Most oligodendrocytes were located at the axonal projections of CA1 neurons located in the stratum oriens. *Right:* Example of a vOL in the hippocampus (cyan, arrow). (G) Left: Schematic of analysis location in the cerebellar Purkinje cell and granule cell layers. *Right:* Example of a vOL in the cerebellum (cyan, arrow). (H) Quantifications of vOLs in the hippocampus and cerebellum (n = 4 mice hippocampus, n = 3 mice cerebellum).
